# Presymptomatic Targeted Circuit Manipulation for Ameliorating Huntington’s Disease Pathogenesis

**DOI:** 10.1101/2024.07.24.604946

**Authors:** Ebenezer C. Ikefuama, Ashley N. Slaviero, Raegan Schalau, Madison Gott, Maya O. Tree, Gary L. Dunbar, Julien Rossignol, Ute Hochgeschwender

## Abstract

Early stages of Huntington’s disease (HD) before the onset of motor and cognitive symptoms are characterized by imbalanced excitatory and inhibitory output from the cortex to striatal and subcortical structures. The window before the onset of symptoms presents an opportunity to adjust the firing rate within microcircuits with the goal of restoring the impaired E/I balance, thereby preventing or slowing down disease progression. Here, we investigated the effect of presymptomatic cell-type specific manipulation of activity of pyramidal neurons and parvalbumin interneurons in the M1 motor cortex on disease progression in the R6/2 HD mouse model. Our results show that dampening excitation of Emx1 pyramidal neurons or increasing activity of parvalbumin interneurons once daily for 3 weeks during the pre-symptomatic phase alleviated HD-related motor coordination dysfunction. Cell-type-specific modulation to normalize the net output of the cortex is a potential therapeutic avenue for HD and other neurodegenerative disorders.

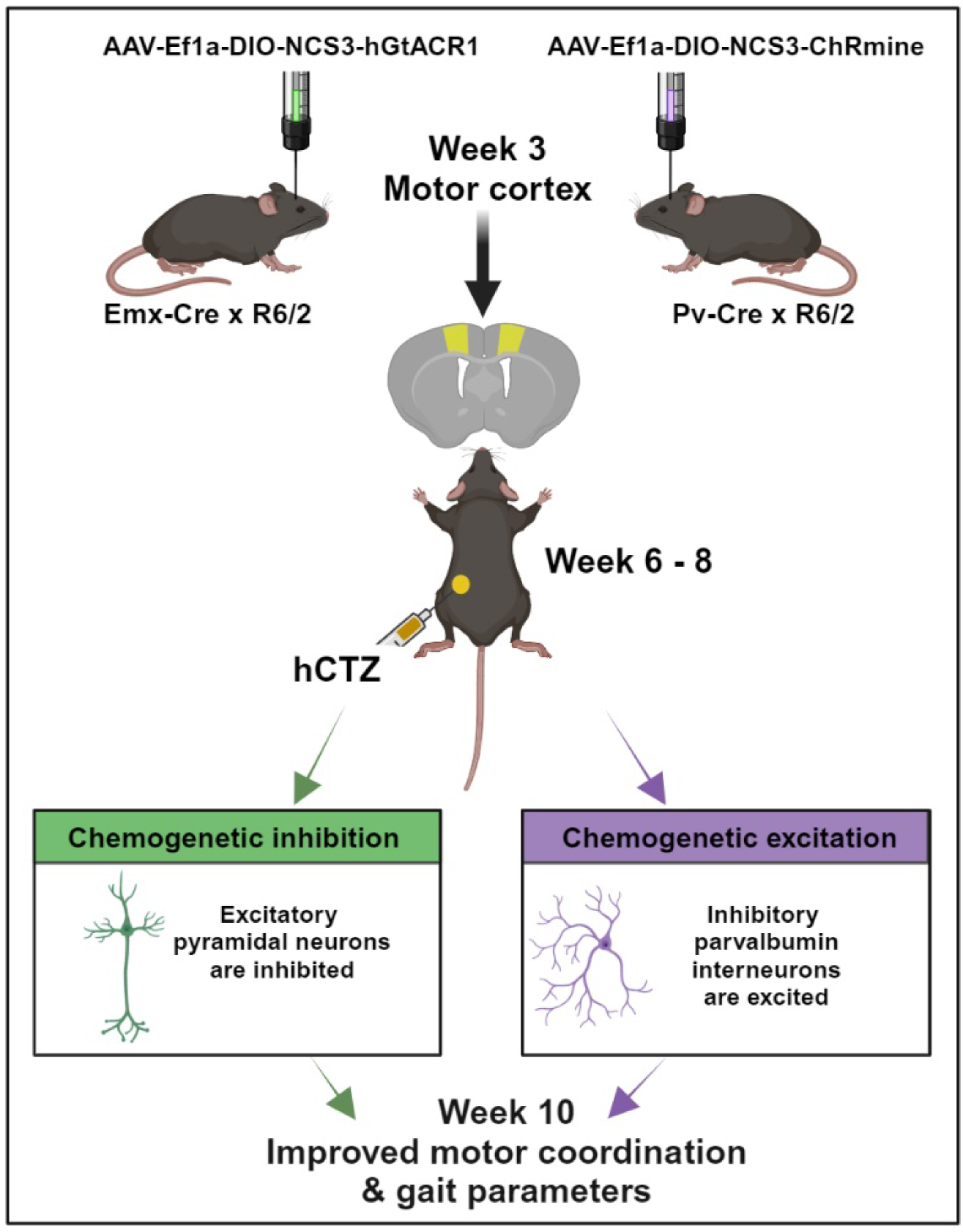

**Highlights:** Cortical excitatory pyramidal neurons and inhibitory parvalbumin interneurons are affected in Huntington’s disease

Repeated bioluminescence-mediated opto-chemogenetic inhibition/excitation of affected cell types in the motor cortex improved motor coordination and inter-limb gait parameters in HD mice

Early manipulation of select microcircuits before the onset of symptoms presents an avenue to slow HD disease progression

## Introduction

Huntington’s disease (HD) is a progressively debilitating neurodegenerative disorder whose prevalence has risen over the past half-century in North America, Australia, and Western Europe, now affecting 1 in 10,000 individuals ^1,2^. The cortico-basal ganglia-thalamo-cortical (CBGTC) loop crucially upholds excitatory/inhibitory (E/I) balance in the normal brain. In postmortem HD brains and genetic HD mouse models, loss of indirect spiny projection neurons (iSPNs), a vital microcircuit within the CBGTC loop, has been identified. This loop encompasses projection neurons (CPNs) in the cortex and subcortical structures—striatum, globus pallidus, subthalamic nucleus (STN), thalamus, and substantia nigra pars reticulata (SNr). The loss of iSPNs disrupts the indirect pathway responsible for repressing globus pallidus external (GPe) neuronal activity. Consequently, this imbalance tilts the network towards continuous and involuntary excitatory states, underlying the hyperactive motor symptoms in early HD stages ^3,4^. This leads to amplified movement initiation (a core role of the direct pathway), shifting the E/I equilibrium toward heightened thalamic and cortical firing states. This hyperexcitability connects to increased direct spiny projection neuron (dSPN) activation and reduced inhibition of thalamic target neurons ^5^.

Studies in rodent HD models during pre- and early symptomatic stages have revealed intricate alterations in intrinsic and synaptic properties of affected neurons. In freely behaving R6/2 transgenic mice, Walker et al. observed elevated spontaneous firing rates, diminished bursting rates, and lost spike synchrony in cortical neurons ^6^. Other *in vivo* electrophysiological examinations demonstrated augmented firing rates in spiny projection neurons during the presymptomatic HD stage ^7,8^. Brain slice whole-cell patch-clamp electrophysiology in BACHD mice exhibited functional shifts, like reduced inhibition onto cortical pyramidal cells and decreased excitation onto parvalbumin (PV) interneurons and pyramidal cells in the same region ^9^.

However, there are currently no effective therapies targeting HD’s underlying pathophysiology. Over two decades post-identification of HD-causing genetic mutations, treatment remains symptom-focused ^10^. Addressing the excitatory and inhibitory imbalance in early HD stages could mitigate a range of symptoms spanning mental health, behavior, movement, and communication.

Deep brain stimulation (DBS) has been explored to restore E/I balance and alleviate motor and cognitive deficits in HD. Yet, this approach often triggers bradykinesia, involuntary muscle contraction, and worsened cognition ^11,12^. DBS’s limitations arise from its non-specific activation of all nearby cells, undermining the intended specificity. Moreover, its invasiveness risks infection, glial scar formation, and electrode degradation_13,14._

On the other hand, optogenetics employs light for precise activation of light-sensing ion channels targeted to genetically defined neural sub-populations ^15^. Optogenetic inhibition of PV-positive neurons in GPe was shown to alleviate hypoactivity in the STN of Q175 mice ^16^. However, behavioral function post-manipulation was not evaluated. Another study demonstrated increased rearing in the open field test and latency to fall during the rotarod test when symptomatic R6/1 mice were given optogenetic excitation of secondary motor (M2) cortical neurons projecting to dorsolateral striatum (M2-DLS) ^17^. However, optogenetics shares with DBS the drawbacks of hardware invasiveness in addition to potential thermal damage from optical fibers ^18^.

Our study aimed to normalize E/I imbalance via cell-type-specific modulation to impede HD progression in mice using Bioluminescent Optogenetics (BL-OG) [ref Berglund 2013, 2016; Medendorp 2021; Petersen 2022; Ikefuama 2022]. This opto-chemogenetic approach leverages biological light generated by a luciferase enzyme upon application of the small molecule luciferin to activate tethered excitatory or inhibitory light-sensing ion channels. BL-OG allows efficient non-invasive manipulation of neuronal subpopulations while addressing limitations of DBS and conventional optogenetics. We demonstrate that reducing cortical drive on the striatum either through direct inhibition of pyramidal neuron activity or through strengthening inhibitory activity of parvalbumin interneurons during the pre- and early symptomatic window of HD ameliorates motor impairment.

## Results

### Presymptomatic circuit manipulation in R6/2 mice

Studies in rodent HD models and in patients have revealed alterations in intrinsic and synaptic properties of cortical neurons during early symptomatic stages ^6,19^. Both increased activity of pyramidal (PYR) neurons and their decreased inhibition by weakened parvalbumin (PV) interneurons result in overall increased drive of cortical output onto striatal neurons ^9,19,20^ (Figure 1A). To counteract the cortico-striatal overdrive before symptoms fully develop, we wanted to express genetically targeted actuators that inhibit the overactive PYR neurons and activate the dampened PV interneurons. We selected an inhibitory luminopsin (LMO) for expression in target 1 (PYR neurons; Figure 1B) and an excitatory LMO for target 2 (PV interneurons; Figure 1C). We chose R6/2 mice for testing the effects of presymptomatic circuit manipulation on HD phenotype. R6/2 mice have a short time span of disease, with no symptoms observed at 5 weeks of age to the full-blown phenotype by 10 weeks; most animals succumb to the disease by 12 weeks. To allow for full expression of AAV-transduced LMOs by 5 weeks of age, we determined the stereotaxic injection coordinates for primary motor cortex (M1) in 3-week-old R6/2 mice using DiI injections followed by fluorescent imaging (Figure 1D). We then transduced Cre-dependent virus into the M1 motor cortex of 3-week-old mice double transgenic for R6/2 and Emx1-Cre (AAV-EfIa-DIO-NCS3-hGtACR1; Figure 1E) or PV-Cre (AAV-EfIa-DIO-NCS3-ChRmine; Figure 1F). Figure 1G outlines the experimental design along a timeline. Viral injections were performed on postnatal day 21 mice, and baseline behavioral tests (rotarod, open-field) were done two weeks later at week 5. This was immediately followed by daily intraperitoneal administration of luciferin (h-coelenterazine, hCTZ) for 3 weeks. At week 10, we performed post-treatment behavioral tests (rotarod, open-field, CatWalk, passive-avoidance) to evaluate the effect of manipulating the activities of cortical PYR neurons and PV interneurons in the M1 cortex. IVIS imaging was performed after the last behavioral test, and mice were perfused and brains evaluated for LMO expression. The weight of these mice was monitored weekly between postnatal day 35 and 70.

**Figure 1.**
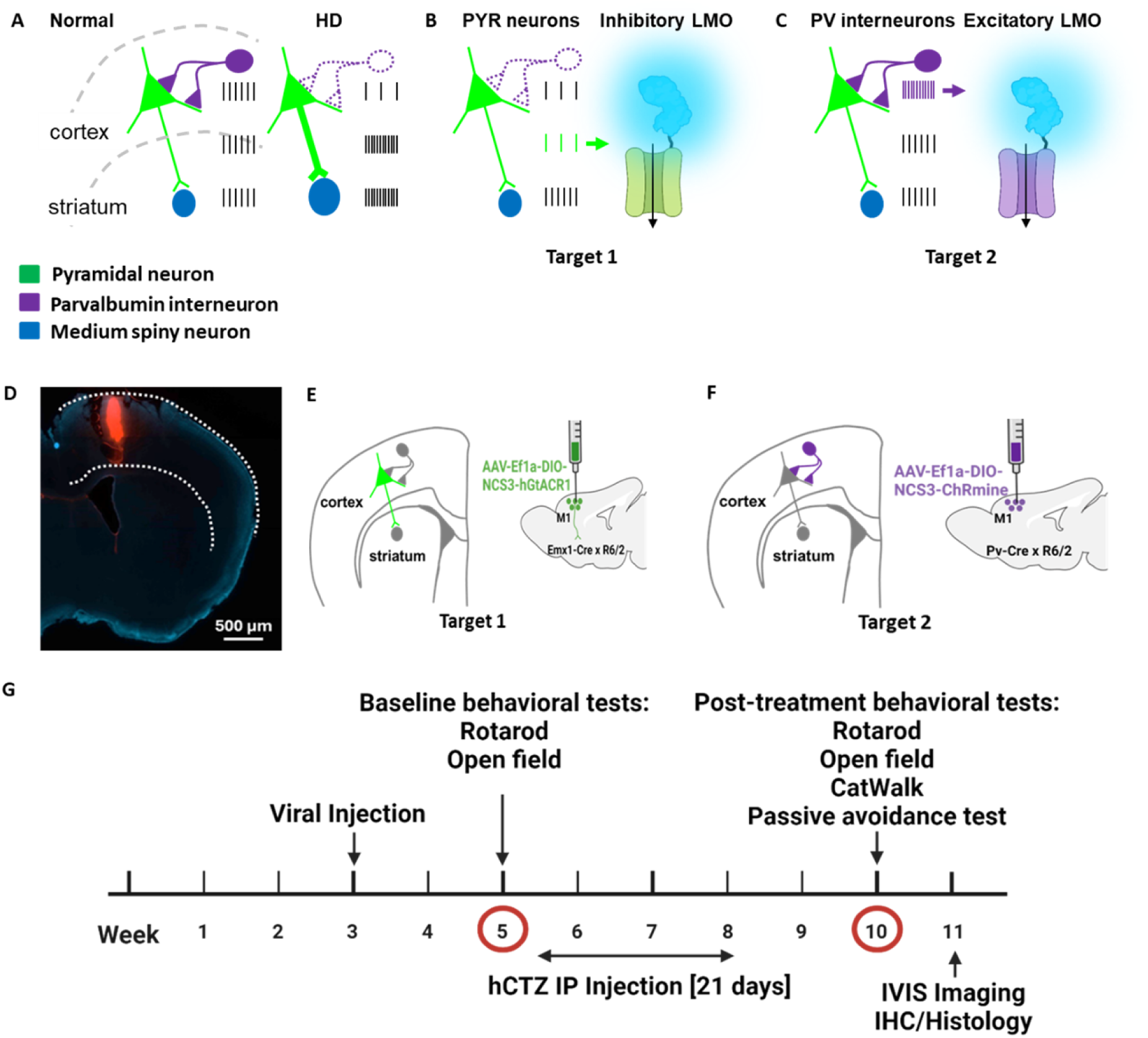
Experimental design for presymptomatic targeted circuit manipulation in Huntington’s disease. (A) Firing patterns of cortical pyramidal neurons (green) and parvalbumin interneurons (purple) in normal compared to hyper- and hypo-active states in HD resulting in increased cortico-striatal drive. (B-C) Normalization of cortico-striatal drive through dampening of pyramidal neuronal activity by expressing an inhibitory luminopsin (LMO) (B) and through increasing parvalbumin interneuron activity by expressing an excitatory luminopsin (LMO) (C). (D) Validation of M1 coordinates in a 3-week-old mouse by Dil injection. (E-F) Neuronal subpopulation-specific targeting of LMOs in M1 cortex by transduction of AAV-Ef1a-DIO-NCS3-hGtACR1 in Emx-Cre x R6/2 mice (E) and AAV-Ef1a-DIO-NCS3-ChRmine in Pv-Cre x R6/2 mice (F). (G) Experimental timeline showing viral injection at week 3, baseline behavioral test at week 5, IP administration of hCTZ from weeks 6 – 8, and post-treatment behavioral tests at week 10.

Five groups of mice, each consisting of males and females, were compared in this study: wildtype mice, untreated R6/2 mice, Emx1-Cre x R6/2 transduced with NCS3-hGtACR1 and treated with hCTZ for 21 days, Pv-Cre x R6/2 transduced with NCS3-ChRmine and treated with hCTZ for 21 days, and R6/2 mice without viral injections but treated with hCTZ for 21 days.

### Efficient circuit manipulation by excitatory and inhibitory luminopsins

For both LMOs, we tethered a bright light emitter, NCS3, to an inhibitory and excitatory opsin. NCS3 is a fusion protein of a molecularly evolved shrimp luciferase, GeNL_SS, and the fluorescent protein mNeonGreen (Lambert et al., 2023; Slaviero et al., 2024), with a peak emission wavelength of 520 nm. For neural inhibition, NCS3 was tethered to hGtACR1, a codon-optimized anion channelrhodopsin 1 from Guillardia theta with a peak activation wavelength of ∼ 540 nm ^23^. For neural excitation we used LMO11, a fusion of NCS3 and ChRmine that showed highly efficient activation by bioluminescence^22^. We validated the functionality of the two LMOs through whole-cell patch clamp recordings in HEK293 cells (Figure 2A-C). LMO expressing cells were stimulated with a green LED for 1 second to determine the maximum photocurrent amplitude. Luciferin (hCTZ) application resulted in robust bioluminescence-induced currents (Figure 2A and B). The coupling efficiencies (hCTZ-induced current amplitude / maximum photocurrent amplitude) of NCS3-GtACR1 and NCS3-ChRmine are 56% and 71%, respectively (Figure 2C).

**Figure 2.**
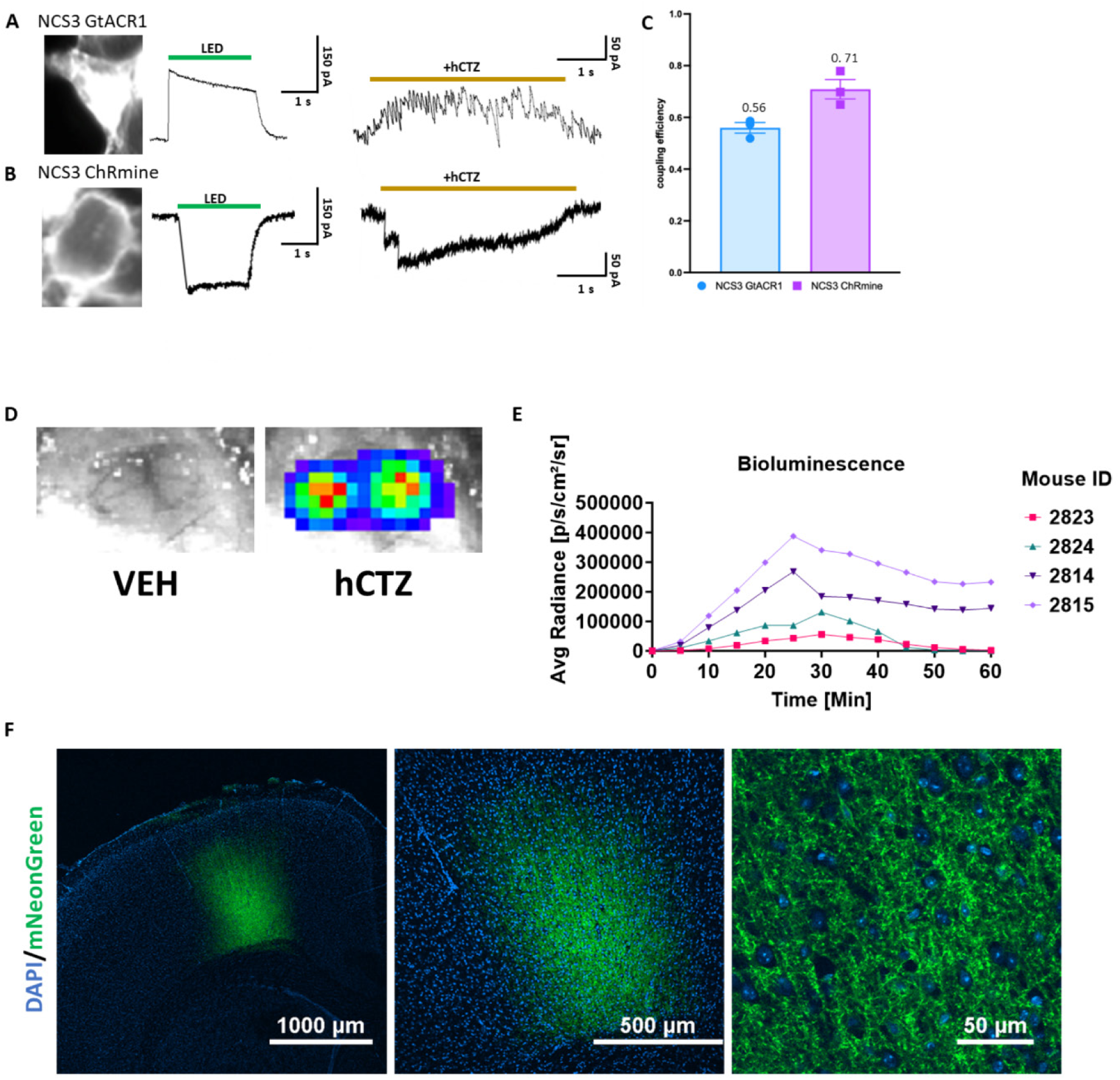
Efficient circuit manipulation by excitatory and inhibitory luminopsins. (A-B) Fluorescent images of LMO-expressing HEK293 cells (left), photocurrent patch clamp traces (middle), and luciferin-induced patch clamp traces (right) for NCS3 – GtACR1 (A) and NCS3 – ChRmine (B). (C) Coupling efficiency of NCS3 - GtACR1 and NCS3 – ChRmine (n = 3/group). (D) Bioluminescence in vivo signal imaged through an open skull from a mouse bilaterally transduced with AAV-EfIa-DIO-NCS3-hGtACR1 following IP delivery of vehicle (left) and hCTZ (right). Exposure time was 1 sec. (E) Bioluminescence emission over time. Images were taken for 1 sec every 5 minutes for 60 minutes after IP injection for 4 individual mice transduced with NCS3-GtACR1 (mouse 2824 and 2814) and NCS3-ChRmine (mouse 2823 and 2815). (F) Fluorescence confocal images of cortical sections showing robust LMO expression (NCS3-ChRmine) at 4x, 10x, and 60x objectives with magnifications as indicated (DAPI = blue, mNeonGreen = green).

To determine the kinetics of LMO activation we imaged, *in vivo*, bioluminescence emission over one hour after ip injection of luciferin (Figure 2D and 2E). Bioluminescence was easily visualized through a craniotomy (Figure 2E). Images taken every 5 minutes show peak bioluminescence – and thus LMO activation – around 25 - 30 minutes after ip application of the luciferin, with substantial light emission still an hour after luciferin application (Figure 2E). Postmortem immunofluorescent imaging of brain sections confirmed robust expression of LMOs in M1 cortex in all mice (see example in Figure 2F).

### Presymptomatic circuit manipulation attenuates motor deficits in R6/2 mice

Directly decreasing activity of Emx1 PYR neurons as well as increasing activity of PV interneurons and thereby decreasing drive of Emx1 PYR neurons from the M1 motor cortex to the striatum is expected to attenuate locomotor impairments in 10-week-old R6/2 mice. We tested this prediction in two experimental paradigms, motor coordination on the rotarod and gait parameters in the Catwalk.

To assess motor coordination, we tested mice using the rotating rod at a fixed speed. At week 5, we habituated the mice on the device for four consecutive days at a fixed speed (10 revolutions per minute, RPM). On the fifth day, we tested these mice at 5 RPM. We observed no genotype difference and no effect of viral injections on the performance of these mice at week 5 of age (Figure 3), indicating the absence of any motor symptoms at this stage in any group. To determine the effect of intraperitoneal administration of hCTZ for the activation of LMOs expressed in PYR neurons (Emx-Inh-hCTZ) and PV interneurons (Pv-Exc-hCTZ), we retested these mice at week 10 of age. While no difference in performance was observed for wildtype mice at 10 weeks, R6/2 mice that were either untreated or only received the substrate, hCTZ, showed severe impairment in motor coordination, barely staying on the rotating rod for even a few seconds (Figure 3). Manipulation of PYR neurons and PV interneurons significantly increased latency to fall off the rotating rod (p ≤ 0.0001 vs. R6/2 – untreated group; p ≤ 0.0003 vs. R6/2 – hCTZ group; Figure 3). There was no significant difference in performance between R6/2 mice with 21 days of dampening Emx1 PYR neurons versus activating PV interneurons.

**Figure 3.**
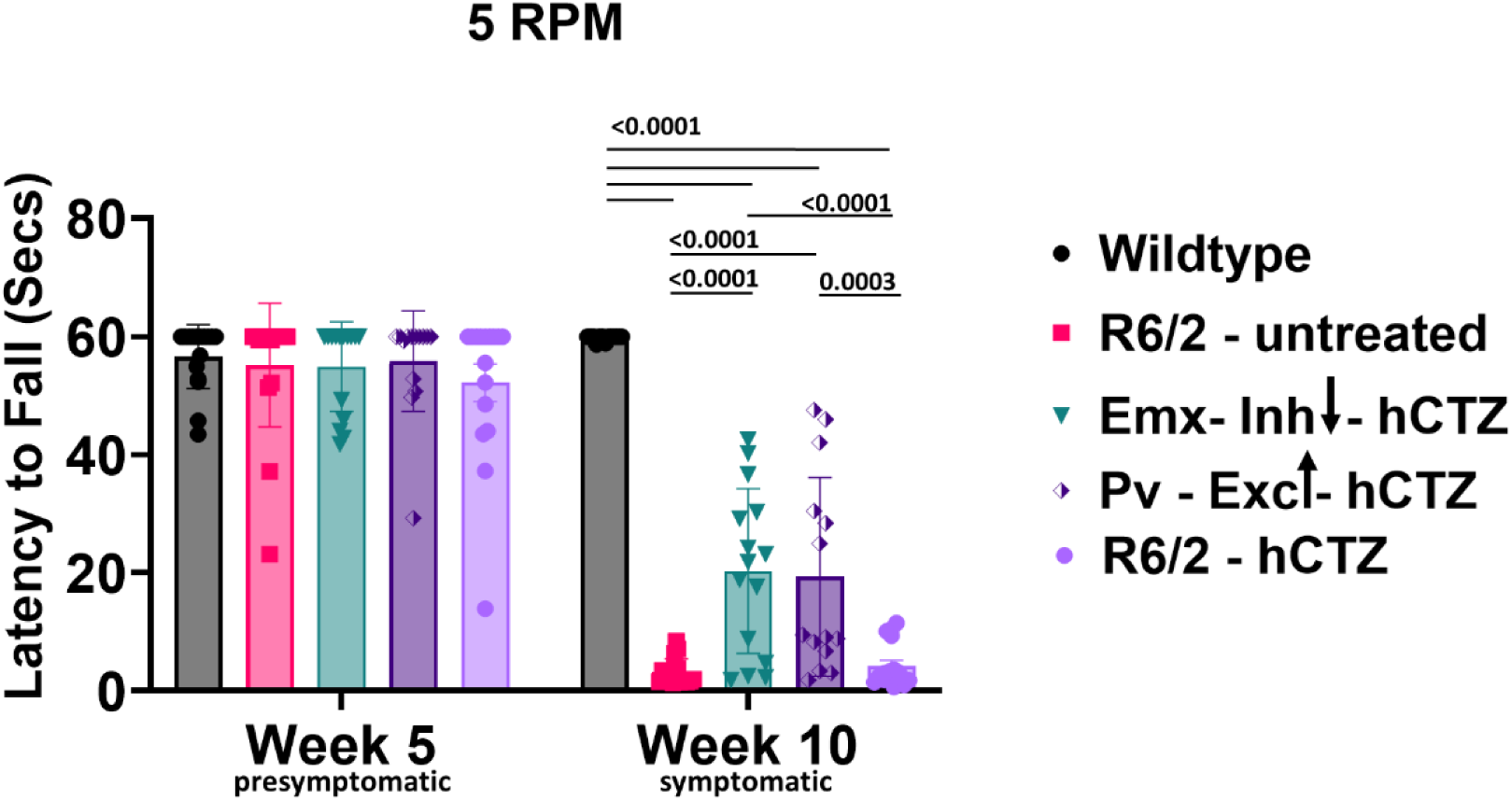
Rescue of motor coordination in R6/2 mice. Average latency to fall from a rotating rod at a fixed speed of 5 revolutions per min. Significance was assessed by two-way ANOVA with Tukey’s post hoc comparison. All data are presented as the mean ± SEM (n = 14 – 16 mice/group).

The Catwalk captures paw placements of freely walking mice to calculate various gait parameters. A comprehensive analysis of 126 parameters revealed significant genotype differences in 46 (Table 2 in the Supplemental Material). Among these, there was a significant treatment effect in 12 parameters (Figure 4 and Supplementary Table 3). These parameters include couplings (Figure 4A), phase dispersions (Figure 4B), and step sequence (Figure 4C). Couplings and phase dispersions, indicative of the temporal relationship between simultaneous paw placements during a step cycle, served as metrics for evaluating inter-paw coordination. Untreated R6/2 mice had paw placements significantly different from wildtype mice (red stippled lines in Figure 4A and B). Manipulating either PYR neurons in Emx-Cre x R6/2 mice or PV interneurons in Pv-Cre x R6/2 mice led to a reduction of deficits observed in untreated R6/2 mice. The rescued limb combinations under couplings and phase dispersions included RF-RH, RH-RF, LF-LH, and LH-LF, implying abnormal right or left limb placement during movement in untreated R6/2 mice (Figure 4A and B). Analysis of the six distinct limb placement sequences disclosed significant genotype and treatment differences in step sequence AB (LH-LF-RH-RF). In this step sequence, there were abnormally high footfall patterns in untreated R6/2 mice, and the inhibition of PYR neurons and excitation of PV interneurons resulted in a significant reduction in the high utilization of this step sequence (Figure 4C). Increase in clasping reflex is one of the early phenotypes of HD progression (Figure 5A) and this is independent of muscle strength and acquired motor skills ^24,25^. There was an absence of clasping behavior in R6/2 mice at week 5. At week 10, all R6/2 mice, irrespective of treatment, showed significant withdrawal of both front and rear paws when compared to wildtype littermates (Figure 5B, p<0.0001) with Pv-Exc-hCTZ mice showing significantly reduced limb clasping compared to untreated R6/2 mice (Figure 5B, p = 0.0230).

**Figure 4.**
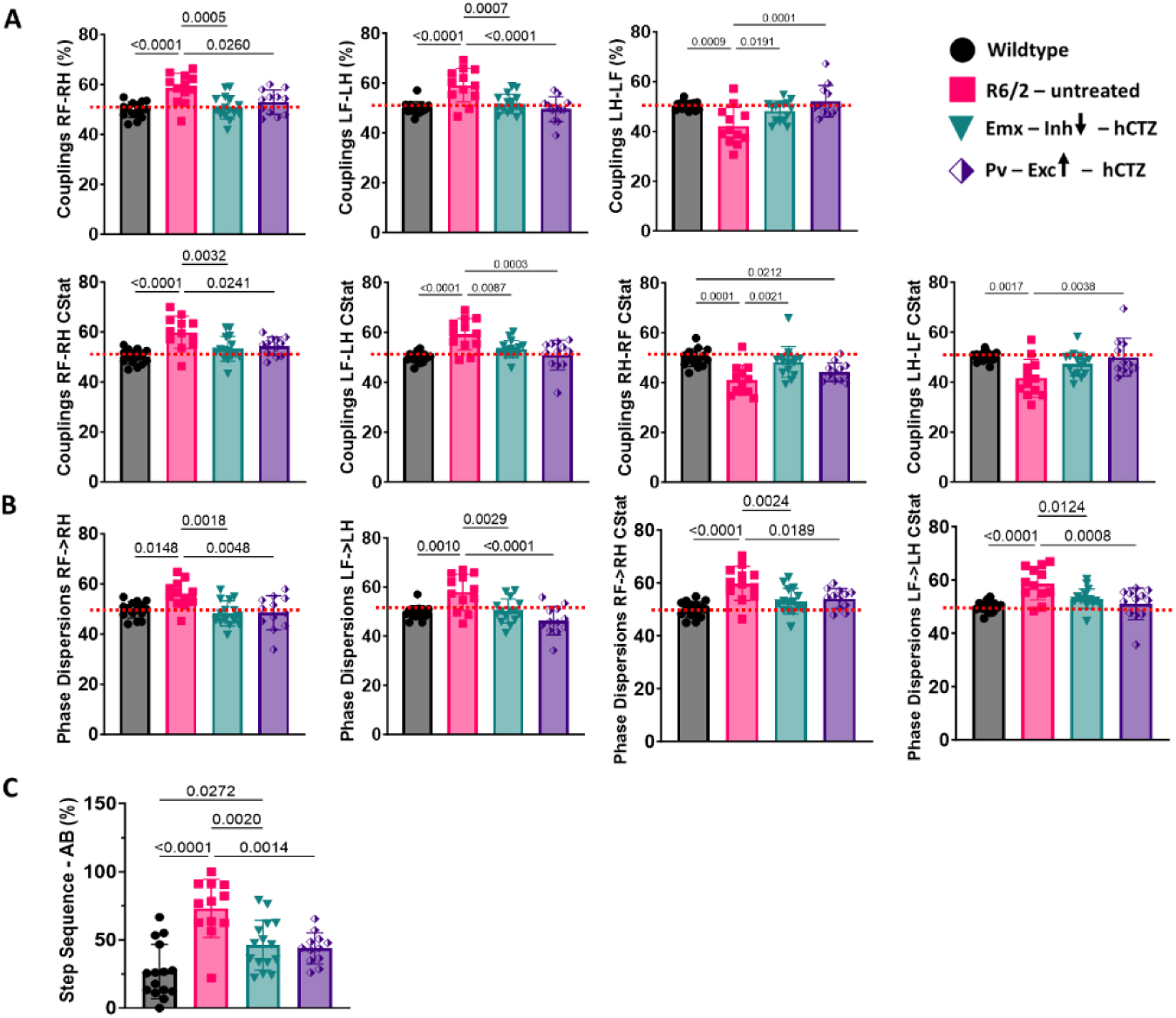
Rescue of gait parameters in R6/2 mice. (A-C) Catwalk data showing significant improvements in Couplings (A), Phase dispersions (B), and Step sequence – AB (C). Significance was assessed by two-way ANOVA with Tukey’s post hoc comparison. All data are presented as the mean ± SEM (n = 12 – 15 mice/group).

**Figure 5.**
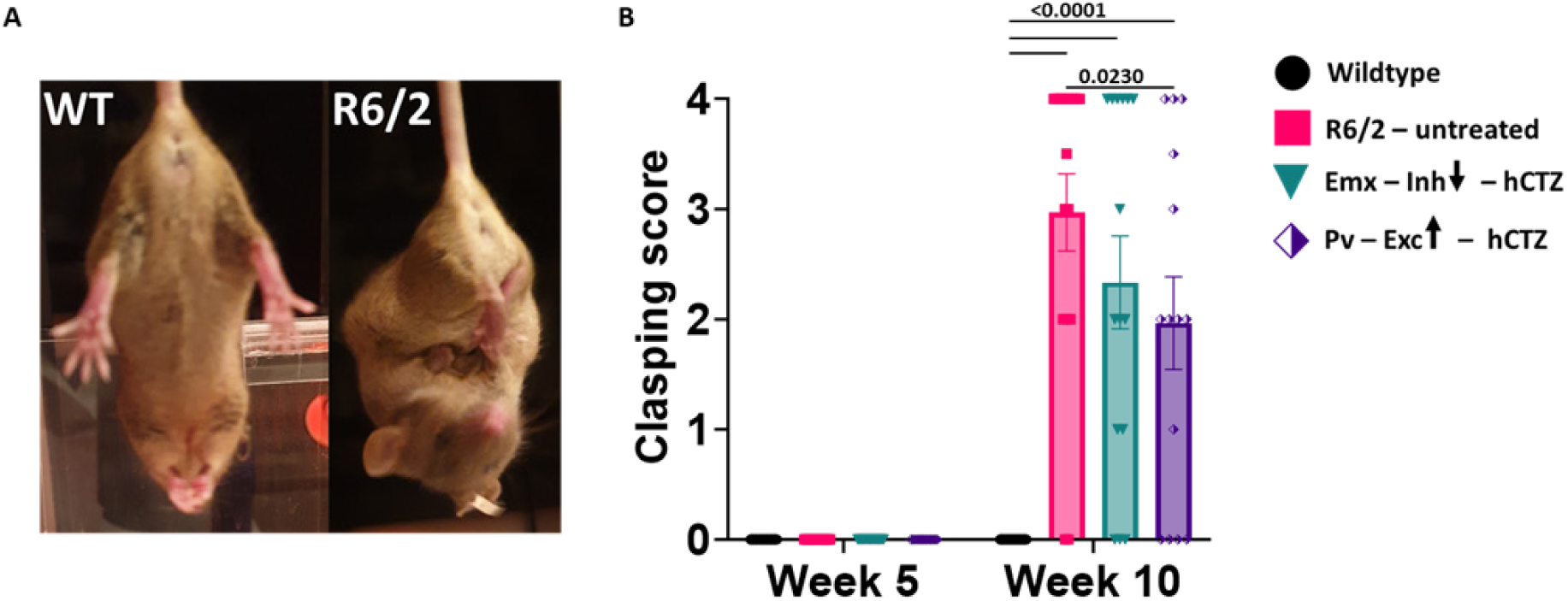
Clasping reflex occurs late in HD pathogenesis in R6/2 mice. (A) Representative images of 10-week-old wildtype and R6/2 mice during a clasping test. (B) Graph shows clasping scores of Wildtype, R6/2 – untreated, Emx – Inh – hCTZ, and Pv – Exc – hCTZ mice at week 5 and week 10. Significance was assessed by two-way ANOVA with Tukey’s post hoc comparison. All data are presented as the mean ± SEM (n = 14 – 16 mice/group).

### Presymptomatic circuit manipulation in motor cortex of R6/2 mice does not affect non-motor phenotypes

To test whether presymptomatic dampening of neuronal drive from the M1 motor cortex to the striatum affects aspects of the HD phenotype other than motor function, we probed exploratory and anxiety behaviors in an open field test and long-term memory impairment in a passive avoidance assay.

Of five parameters for exploratory behavior tested in 5-week-old mice, four showed already significant differences between wildtype and R6/2 mice at this early stage (Figure 6A and 6C). By 10 weeks of age, R6/2 mice were significantly impaired in all categories tested and none of the behaviors improved with our circuit manipulations (Figure 6A-C). R6/2 mice displayed significantly fewer basic (large animal movements from one point to another, p = 0.0007) and fine (small animal movements such as grooming, p < 0.0001) movements. These reductions in movements were not rescued with the manipulation of the two distinct neuronal populations in the M1 cortex (Figure 6A). While rearing behavior (standing on rear limbs) was normal at week 5, R6/2 mice, irrespective of treatment, showed a significant decline on this measure at week 10 (Figure 6B). R6/2 – untreated, Emx-Inh-hCTZ and Pv-Exc-hCTZ groups of mice covered significantly shorter distances in the center of the open field compared to wildtype mice at weeks 5 and 10 (Figure 6C, p = 0.0029 and p = 0.0001, respectively). BL-OG manipulation of pyramidal neurons and parvalbumin interneurons did not alleviate these deficits (p < 0.0001). Distance traveled in the periphery by these mice showed a similar result. Interestingly, all groups of mice spent equal amounts of time in the center and periphery of the grid, suggesting that these mice were not anxious (Figure 6D). Further, there were no differences when tested at 5 weeks versus 10 weeks.

**Figure 6.**
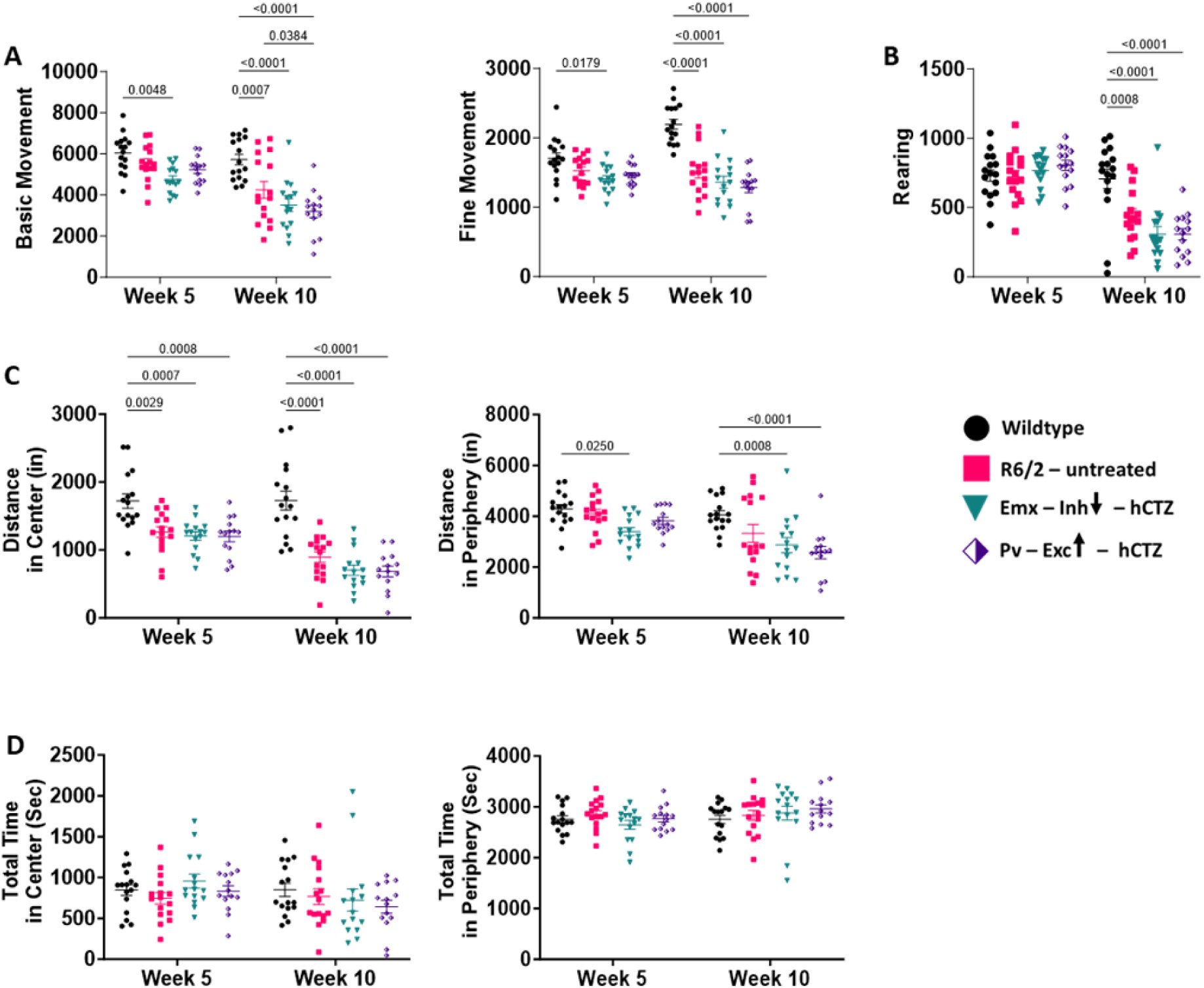
Presymptomatic circuit manipulation in motor cortex of R6/2 mice does not affect exploratory and anxiety-like behavior in R6/2 mice. (A – D) Open field behaviors of wildtype and R6/2 mice. None of the tested parameters were affected by BL-OG stimulation of M1 cortical neurons. Significance was assessed by two-way ANOVA with Tukey’s post hoc comparison. All data are presented as the mean ± SEM (n = 14 – 16 mice/group). (A) R6/2 mice made significantly fewer basic and fine movements starting at week 5 and much more pronounced at week 10 compared to wildtype mice. (B) Rearing behavior in R6/2 mice was significantly affected at week 10, as the disease progressed. (C) R6/2 mice covered less distance in center and periphery starting at week 5 and more pronounced at week 10 compared to wildtype mice. (D) There were no noticeable differences in the total time spent either in the center or periphery of the cage between wildtype and R6/2 mice, either treated or untreated, at either age group.

To measure long-term memory, mice were tested for their ability to remember and avoid entering a shock-associated chamber. At 0 hrs, there were no significant differences between groups. At 24 hours after the administration of footshock, results showed that all R6/2 mice had significantly lower latencies compared to wildtype mice to enter the dark chamber and this tendency was not affected by either the inhibition of pyramidal neurons or the excitation of parvalbumin interneurons of the M1 cortex (p ≤ 0.0001 vs. wildtype group; Figure 7). Within-group comparison showed the expected significant difference between 0 hrs vs 24 hrs in wildtype mice (p < 0.0001), and, interestingly, also in Pv – Exc – hCTZ mice (p = 0.0009), indicating presence of some memory function in this treated group.

**Figure 7.**
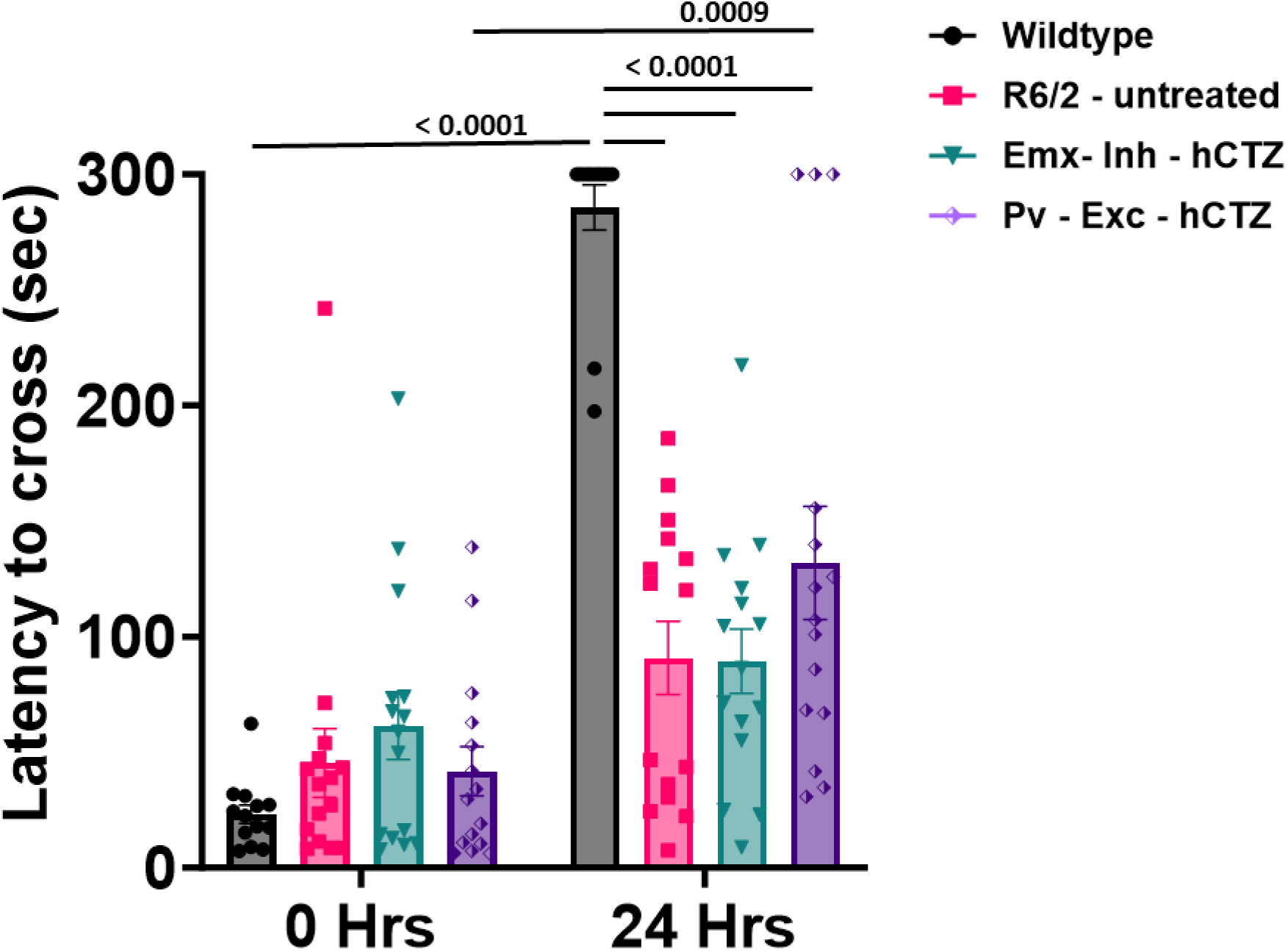
Presymptomatic circuit manipulation in motor cortex of R6/2 mice does not affect memory impairment in R6/2 mice. Memory performance was assessed as the ability to remember a footshock experienced 24 hours before testing. Significance was assessed by two-way ANOVA with Tukey’s post hoc comparison. All data are presented as the mean ± SEM (n = 14 – 16 mice/group).

### Neuronal manipulation showed no effect on body weight

To assess whether circuit manipulation affects overall health of the mice, we recorded the weights of all animals weekly from week 5 until week 10. We observed a steady increase in the weight of female and male mice up to weeks 8 - 9, without any significant differences between genotypes or treatment groups. At week 10, there was a slight decrease in the weight of female R6/2 mice that reached significance compared to wildtype mice only for the Pv-Exc-hCTZ group (Figure 8A). Treated, but not untreated male R6/2 mice showed a significant decrease in weight compared to wildtype at week 9, and all R6/2 males had significantly lower weight than wildtype mice by week 10 (Figure 8B). These findings indicate that our circuit manipulations do not overtly interfere with the animals’ health nor alleviate the overall decline observed in R6/2 mice at later stages.

**Figure 8.**
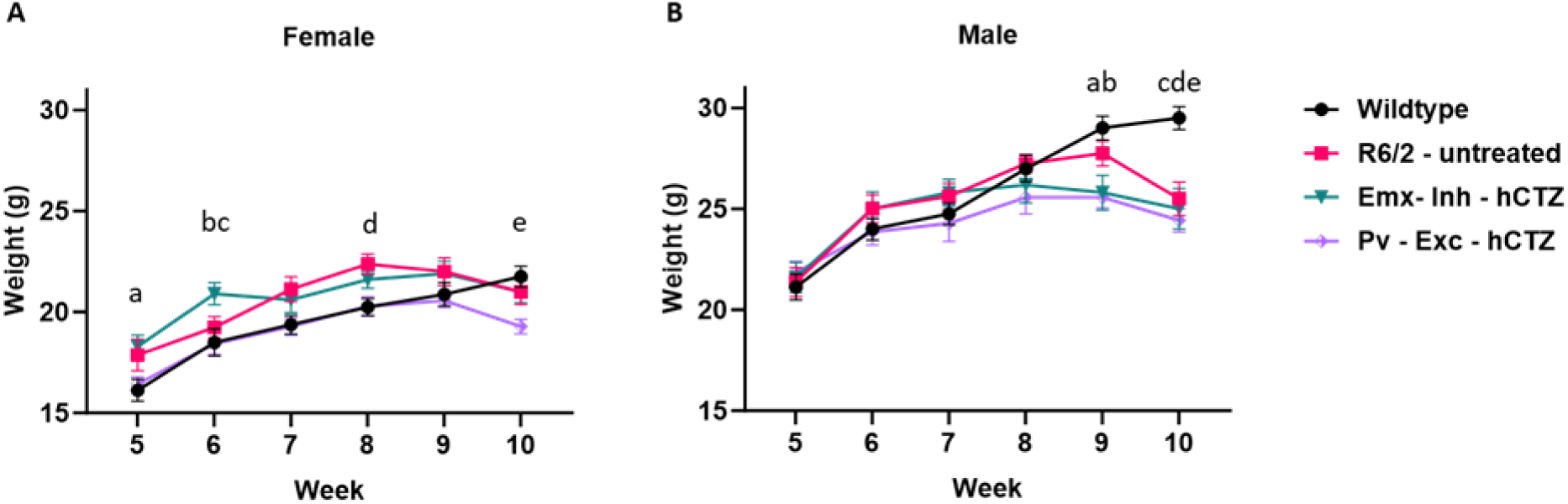
Average weight of wildtype and R6/2 mice across different timepoints. (A) Multiple comparisons of female mice; a = Wildtype vs Pv-Exc-hCTZ, p = 0.0236; b = Wildtype vs Emx-Inh-hCTZ, p = 0.0097; c = Emx-Inh-hCTZ vs Pv-Exc-hCTZ, p = 0.0106; d = Wildtype vs R6/2-untreated, p = 0.0420 and e = Wildtype vs Pv-Exc-hCTZ, p = 0.0171. (B) Multiple comparisons of male mice; a = Wildtype vs Emx-Inh-hCTZ, p = 0.0120; b = Wildtype vs Pv-Exc-hCTZ, p < 0.0019; c = Wildtype vs R6/2-untreated, p = 0.0001; d = Wildtype vs Emx-Inh-hCTZ, p = 0.0001; and e = Wildtype vs Pv-Exc-hCTZ, p < 0.0001. Significance was assessed by two-way ANOVA with Tukey’s post hoc comparison. All data are presented as the mean ± SEM (n = 5 – 10 mice/group).

## Discussion

Multiple morphological and electrophysiological abnormalities of cortical neurons have been discovered in mouse HD models and pathological changes in both patients and mouse models have been observed in the neocortex well before any changes in the striatum ^2,26–29^. Alterations in synaptic activity in the motor cortex observed in patients can be sufficient to produce changes in motor behavior in the absence of striatal dysfunction ^30,31^. Whereas neuronal death may underlie many symptoms in late stage HD^32^, early deficits, which are apparent years before cell death or neurological symptoms, are more likely associated with cellular and synaptic dysfunction in the cortex ^33–37^. This present study was undertaken to determine if chronic stimulation (inhibition/excitation) counteracting these abnormal activities during the presymptomatic stage could attenuate the behavioral deficits seen in later stages of HD. The results demonstrate that either the inhibition of cortical pyramidal neurons or excitation of inhibitory parvalbumin interneurons in M1 cortex does improve motor coordination on the rotarod and inter-limb gait performance on the CatWalk in R6/2 mice. However, the manipulation of the M1 cortex did not rescue memory and exploratory behavior deficits as shown in passive avoidance and open field tests, respectively. Taken together, the data generated from this study reveals that bioluminescent-driven optogenetic modulation of neuronal activities before the onset of symptoms attenuates Huntington’s disease motoric phenotypes.

First, we validated the functionality of the two LMOs used here by *in vitro* patch-clamp recordings of HEK cells expressing NCS3-GtACR1 and NCS3-ChRmine. Application of light led to the expected changes in current amplitude in response to LED and hCTZ, respectively. *In vivo* bioluminescence imaging of LMOs, previously shown to be a proxy of opsin activation ^38,39^, revealed peak light emission and thus neuronal activation/silencing around 25 to 30 minutes after ip injection of the luciferin and lasting for over an hour.

We explored the effects of daily LMO activation over 3 weeks during the presymptomatic phase of HD on end-stage phenotypes at 10 weeks of age. In this study, the primary motor cortex (M1) was targeted because of its role in the control of motor movements that is impaired in HD. To confine expression of inhibitory LMOs to cortical pyramidal neurons and of excitatory LMOs to parvalbumin interneurons, we transduced M1 cortical neurons in Emx-Cre- and Pv-Cre-R6/2 mice, respectively, with Cre-dependent viral vectors.

Baseline behavioral assessments revealed no observable motoric deficits at week 5 confirming the absence of any potential deficits caused by viral injections and establishing a baseline for identifying any treatment effects observed at week 10. Behavioral assessments conducted at week 10 revealed genotype differences in rotarod and CatWalk performances. The manipulation of excitatory pyramidal neurons and inhibitory parvalbumin interneurons both led to an improvement in motor coordination and gait performance. The improvements seen with either target manipulation were comparable, indicating that net excessive drive of M1 output on striatal neurons is a pathogenetic cause of impaired motor performance in HD and a potential therapeutic target.

Not surprisingly, BL-OG-mediated stimulation of cortical pyramidal neurons and parvalbumin interneurons led to no rescue of deficits in non-motor related phenotypes. In the open field task both treated groups of mice showed the same significant decrease in movement, rearing, distance traveled in center and periphery as untreated R6/2 mice. Contrary to other studies, untreated and treated R6/2 mice used in this study showed no anxiety-like phenotype in that they spent equal amount of time in the center and periphery of the open field ^40–42^.

Memory loss is another feature in HD disease progression. Here, we wanted to assess the effect of M1 circuit manipulation on long term memory. We observed no BL-OG-mediated changes in the inability of R6/2 mice to remember a previous footshock. Targeting circuits in the prefrontal cortex, hippocampus, and striatum other than the M1 cortex might yield better results due to their roles in learning and memory consolidation_43–47._

The present study has provided the first direct and specific evidence that targeted manipulation of cortical neurons in the M1 cortex led to the amelioration of motor deficits. Our study introduced a promising approach, BL-OG, for cell-type-specific modulation in HD. This method offers potential for therapeutic advancement in treating HD and may have broader implications for neurodegenerative disorders characterized by E/I imbalances.

### Limitations of the study

In this study, we tested the effects of dampening excitation of Emx1 pyramidal neurons or increasing activity of parvalbumin interneurons in presymptomatic HD mice over a relatively broad temporal window, weeks 6 - 8. This time window is sufficient to observe amelioration of motor deficits. It will be interesting in future studies to decrease the treatment time and to shift the treatment window across the lifespan of the animal.

In this initial investigation we chose to counteract the pathological activity of pyramidal neurons and of parvalbumin interneurons, based on observations in patients and HD animal models. This does not mean that other cell types in the cortex are not involved or are not amenable to activity modulation with beneficial effects. The role of other neuronal cell types can be investigated taking advantage of the many available cell-type specific Cre driver lines.

With our behavioral testing we focused on the assessment of key motor skills and some aspects of “higher order” and cognitive function. Our goal was to identify robust effects of presymptomatic activity modulation rather than exhaustive behavioral testing which, again, will be interesting to examine in future studies.

Our study is using the R6/2 mouse model of HD. We are well aware of the multiple other mouse models of HD, all of them with their specific strengths and weaknesses. It will be interesting to see presymptomatic neural activity interventions applied in several other HD mouse models.

In this study, we found that counteracting the imbalanced excitatory and inhibitory output from the cortex to striatal and subcortical structures during a presymptomatic window lessens motor deficits in HD mice. However, the underlying mechanisms remain unknown. Future work should be directed towards elucidating the functional connections between cortical projection neurons and striatal neurons and their alterations by circuit manipulations.

## Acknowledgements

We thank all members of the Bioluminescence Hub laboratories (http://www.bioluminescencehub.org/) for their feedback, discussions, and thoughtful comments throughout these studies. This work was supported by the National Institutes of Health (R21NS132089, U01NS099709) and the National Science Foundation (NeuroNex-1707352).

## Author contributions

Conceptualization, U.H. and E.C.I.; Methodology, E.C.I., G.L.D., J.R. and U.H.; Formal Analysis, E.C.I., A.N.S., A.D.S. and E.L.C.; Investigation, E.C.I., A.N.S., A.D.S., R.S., M.G. and M.O.T.; Resources, G.L.D., J.R. and U.H.; Writing – Original Draft, E.C.I.; Writing – Review & Editing, E.C.I., G.L.D., J.R. and U.H.; Supervision, G.L.D., J.R. and U.H.; Funding Acquisition, J.R. and U.H.

## STAR Methods

### Lead contact

Further information and requests for resources and reagents should be directed to and will be fulfilled by the lead contact, Ute Hochgeschwender.

### Materials availability

This study did not generate new unique reagents.

### In vitro electrophysiology

Cell culture of human embryonic kidney fibroblasts (HEK293, RRID:CVCL_0045) and electrophysiological recordings were performed following the previously described methodology (Slaviero et al., 2024). Lipofected HEK293 cells were plated sparsely on 15 mm coverslips (Neuvitro) coated with poly-D-lysine in a 12-well plate. Electrophysiological recordings were performed 24-72 hours after lipofection. The recording chamber, containing a coverslip, was continuously perfused with ACSF at a flow rate of 1.5 mL/min. The ACSF solution contained specific concentrations of NaCl, D-glucose, HEPES, KCl, CaCl_2_, and MgCl_2_, which were maintained at 150 mM, 20 mM, 10 mM, 3 mM, 2 mM, and 2 mM, respectively. The pH of the solution was kept at 7.4, the osmolarity was approximately 300 - 315 mOsm/kg, and the temperature was maintained at 35 ± 1°C. The intracellular solution utilized contained potassium gluconate, HEPES, potassium chloride, sodium phosphocreatine, sodium ATP, and sodium GTP, maintained at concentrations of 130 mM, 15 mM, 8 mM, 5 mM, 4 mM, and 0.3 mM, respectively. The pH was approximately 7.25 and the osmolality was around 295 - 300 mOsm/kg. Borosilicate glass micropipettes with an average resistance of 4 MΩ were fabricated using a vertical puller (PC-100, Narishige). Clusters of 5‒15 cells expressing mNeonGreen were visually recognized under a fluorescence microscope. A halogen light source (130 W, U-HGLGPS, Olympus) and filter cubes designed for green excitation (Ex/Em: 540/600, U-MWIGA3, Olympus) were used to stimulate ChRmine and GtACR1. Based on the peak bioluminescent emission of the light emitter, photocurrents were produced by a 1-second exposure to green light. A maximum light intensity of 36.5 mW/cm^2^ irradiance was recorded using a light meter (ThorLabs). Utilizing a Digidata 1440 digitizer, Multiclamp 700b amplifier, and pClamp 10 recording software (Molecular Devices, San Jose, CA, USA)), whole-cell voltage-clamp recordings were performed at -60 mV. Following cell break-in, a gap-free technique was used to assess the photocurrent response, and hCTZ was then applied by perfusion. A new stock of hCTZ (50 mM in NanoFuel) was diluted to 500 µM in ACSF, resulting in a total of 500 µL. For each recording, fresh hCTZ working stock was prepared and added right away to the perfusion chamber (final concentration in the bath was around 100 µM). The recordings were collected at 10 kHz and filtered via a Gaussian lowpass filter with a 250 Hz -3dB cutoff on Clampfit. Photocurrents were assessed during the last 100 milliseconds of illumination, specifically at the steady state level. This was done to eliminate any irregularities noticed before light adaption, which is when the strongest current response occurs. The Luciferin-induced currents were measured and analyzed using Clampfit software. The amplitude of the currents reached a consistent level once the stimulation was applied. The coupling efficiencies were calculated by normalizing the change in current produced by luciferin to the change in photocurrent elicited by the lamp, specifically by dividing the luciferin-induced current by the photocurrent.

### Animals

All experimental procedures were conducted in accordance with standard ethical guidelines and were approved by Central Michigan University Institutional Animal Care and Use Committee (IACUC ID # 2020-1007 and #2023-733). All animals were maintained on a 12 h/12 h reverse light-dark cycle (with lights on/off at 12 A.M./ 12 P.M.) at constant temperature (22 °C) and humidity (55±15 %), with free access to food and water. All in vivo studies were performed on wildtype, R6/2, Emx-Cre x R6/2 and Pv-Cre x R6/2 mice generated by crossing wildtype females transplanted with ovaries from transgenic R6/2 female mice (B6CBA-Tg(HDexon1)62Gpb/1J; Jackson Laboratory, Stock No. 002810) with wildtype (B6CBA), Emx-Cre (Jackson #005628), and Pv-Cre (Jackson #017320) males.

### Genotyping and random animal assignment

Toe snips were collected at postnatal day 7 and sent to the Genotyping Center of America (GTCA) to perform genotyping for R6/2, Emx-Cre and Pv-Cre via PCR analysis using genomic DNA. On postnatal day (PD) 21, experimental animals were weaned from multiple litters. To limit the effects of litter origin on the experimental results, two to four mice were allocated from the same litter into the same experimental group ^48,49^. Supplementary Table 1 lists the distribution of mice across sex, genotypes, and litters. Mice were housed in social groups with same-gender littermates in cages containing bedding and nesting material.

### Stereotaxic surgery and viral transduction

Stereotactic cranial window surgery and intracortical delivery of adeno-associated virus (AAV) was performed on three-week-old R6/2 x Emx-Cre and R6/2 x Pv-Cre mice under isoflurane at a rate maintained between 2 – 3% and oxygen at 55 mL/min (SomnoSuite Low-Flow Anesthesia System, Kent Scientific Corporation). A 10 μL World Precision Instruments syringe with 35G beveled needle was lowered into the brain with coordinates 1 mm (AP), ±1.3 mm (ML) and -1 mm (DV) with respect to bregma for injections into the M1 cortex of both hemispheres. One microliter was injected at a rate of 200 nL/min of AAV-Ef1a-DIO-NCS3-hGtACR1 (VectorBuilder, titer: 1.3 x 1013) into R6/2 x Emx-Cre mice (Target 1) and AAV-Ef1a-DIO-NCS3-ChRmine (VectorBuilder, titer: 6.56 x 1012) in R6/2 x Pv-Cre mice (Target 2).

### Intraperitoneal administration of h-Coelenterazine (hCTZ)

Lyophilized h-Coelenterazine (NanoLight Technology, CAT#3011) was dissolved in sterile water to a concentration of 1µg/µL and 200 µL of the solvent was administered intraperitoneally to each mouse daily for 21 days.

### Behavioral assessments

All behavioral assessment procedures were performed during the animal’s dark cycle.

### Rotarod

Motor coordination was assessed by measuring the latency to fall from a San Diego Instruments Rotarod (San Diego, CA). In week 5, mice were habituated on days 1 – 4 at 10 RPM for 2 minutes daily and tested at 5 revolutions per minute (RPM) on day 5. At week 10, habituation was omitted, and mice were tested at 5 RPM.

### CatWalk quantitative gait analysis test

CatWalk XT 9 (Noldus, The Netherlands) was utilized to evaluate gait and locomotion. The animals freely explored a green illuminated glass plate, and a high-speed video camera captured the reflected light from paw contacts. A 9 mm recording section with automatic detection settings was employed, utilizing a green intensity threshold of 0.1, ceiling light of 17.7 V, and a camera gain of 20 dB. Testing occurred in darkness, and the animals voluntarily performed back-and-forth runs. The three fastest trials were selected and averaged for subsequent analysis ^50^.

### Open field (OF) test

Exploratory and anxiety-like behavior was assessed in an open field box. The OF testing setup comprised a Plexiglas enclosure measuring 47.5 cm x 25.5 cm x 21 cm. It was equipped with a Motor Monitor, Version 1.2, from Hamilton-Kinder (Chula Vista, CA), featuring grids of infrared beams positioned 2.5 cm above the OF floor for assessing horizontal movement and 7.5 cm above the OF floor for measuring vertical activity, such as rearing, along the perimeters of the OF. The infrared grids were comprised of 16 photobeams in both the horizontal and vertical directions, resulting in a total of 16x16 photobeams. These grids allowed for the tracking of the mouse’s location by detecting when the infrared beams in the area were obstructed by the mouse’s motions. The software was linked to the device utilized for quantifying the mice’s total locomotion, as determined by the frequency of interruptions in the gridded infrared beam system. The OF test was recorded for one hour and the mice were carefully taken out of the box and placed back into their home cages.

### Passive avoidance

The passive-avoidance (PA) test consisted of two separate days: the first day for "training" and the second day for "testing," with a 24-hour gap between them. During the training session, each mouse was placed in the well-lit section of a PA apparatus (Maze Engineers, Skokie, IL) with the connecting door closed. Following a 30-second habituation period, the time taken by each mouse to enter the dark compartment was recorded to evaluate potential differences in the overall motivation to explore this unfamiliar environment based on genotype. As the mouse entered the dark compartment, the guillotine door closed, and it received two footshocks immediately after entering the dark compartment. Each footshock had a strength of 0.3 mA and lasted for 3 seconds, with a 1-second delay between shocks. After the test, the mice were left in the dark chamber for a duration of 10 seconds, after which they were taken out and returned to their home cages. Memory retention was assessed by recording the duration it took for the mice to enter the dark chamber 24 hours following the footshock administration ^51^.

### Clasping

An increase in the clasping of limbs is associated with disease severity and progression. Here, we tested all mice by holding them by their tail for 30 seconds and counted the number of clasped limbs. We assigned a score of 0, 1, 2, 3 or 4 to indicate the absence or presence of clasping behavior of one, two, three or four limbs, respectively. The average of two trials was used for further analysis ^52^.

### IVIS imaging

Bioluminescence in mice was assessed using an IVIS Lumina LT in vivo bioluminescence imaging system (Perkin Elmer). Upon conclusion of the behavioral assessment, anesthetized mice received hCTZ intraperitoneally and were immediately placed inside the imaging chamber. Bioluminescence images were captured using the specified parameters: binning set to Large, F/Stop at 1, exposure duration of 1 minute, emission filter set to open, excitation filter blocked, and a field of view (FOV) of 5 cm (A). Regions of interest (ROIs) were measured for each image to obtain values for the average radiance (p/s/cm^2^/sr) for each mouse.

### Brain tissue preparation and confocal microscopy

After IVIS imaging, 11-week-old mice were anesthetized by isoflurane and transcardially perfused with 1x PBS followed by 4% paraformaldehyde (PFA). The brains were isolated, placed in 4% PFA and stored at 4 degrees Celsius overnight. After 24 hours, they were transferred to 30% sucrose until they sank. Brains were flash frozen in 2-Methylbutane (chilled on dry ice) and embedded in M-1 Embedding Matrix (Ephredia). They were cut at 50 μm using Cryostar NX50 V Low Profile cryostat (Thermo Fisher Scientific) and sections were mounted onto gelatin-coated slides. Images were acquired using 4x (air), 10x (air), and 60x (water) objectives under a Nikon A1R laser scanning confocal microscope. We chose the appropriate channels and laser power to capture mNeonGreen and DAPI fluorescence, and NIS Elements AR was used to process the images further.

## Data analysis

Data were analyzed for statistical significance using GraphPad Prism software version 10. Statistical tests involved either one- or two-way ANOVA, followed by Tukey’s post-hoc test to identify significant interactions. A p-value less than 0.05 was considered statistically significant. Specific details of the statistical tests are provided in the figure legends, including the mean ± SEM for each experiment. For behavioral experiments, groups included a minimum of 12 mice each, ensuring enough statistical power to detect effects at a p-value of 0.05. We assessed for sex differences in the rotarod test and found no statistically significant differences between males and females. To ensure statistical power and to simplify the presentation of our findings, we combined data from both sexes for subsequent analyses.

## Supplemental Information

Table S1. Random and unbiased distribution of mice for all behavioral experiments, related to STAR methods.

Table S2. 46 CatWalk gait parameters showing genotype differences for R6/2 mice, related to Figure 4.

Table S3. Two-way ANOVA with Tukey’s post-hoc comparison of gait parameters showing significant genotype and treatment differences, related to Figure 4.

