## Supplementary Table 1 for "Presymptomatic Targeted Circuit Manipulation for Ameliorating Huntington’s Disease Pathogenesis"

### **Table S1: Random and unbiased distribution of mice for all behavioral experiments**

| **Group** | **Number of mice** | | **# of litters** |
| --- | --- | --- | --- |
|  | **Male** | **Female** |  |
| **Wild-type** | **8** | **8** | **10** |
| **R6/2 - untreated** | **8** | **8** | **13** |
| **R6/2 - Emx - hCTZ** | **5** | **10** | **7** |
| **R6/2 - Pv - hCTZ** | **7** | **7** | **6** |
| **R6/2 - hCTZ** | **7** | **7** | **8** |
